# Too little and too much: balanced hippocampal, but not medial prefrontal, neural activity is required for intact novel object recognition in rats

**DOI:** 10.1101/2025.04.01.646403

**Authors:** Charlotte J. L. Taylor, Jacco G. Renström, Joanna Loayza, Miriam Gwilt, Stuart A. Williams, Rachel Grasmeder Allen, Paula M. Moran, John Gigg, Joanna Neill, Michael Harte, Tobias Bast

**Affiliations:** School of Psychology and Neuroscience@Nottingham, University of Nottingham, UK, NG7 2RD; Division of Neuroscience and Experimental Psychology, University of Manchester, UK, M13 9PT; Division of Pharmacy and Optometry, University of Manchester, UK, M13 9PT

## Abstract

Impaired GABAergic inhibition, so-called neural disinhibition, in the prefrontal cortex and hippocampus has been linked to cognitive deficits. The novel object recognition (NOR) task has been used widely to study cognitive deficits in rodents. However, the contribution of prefrontal cortex and hippocampal GABAergic inhibition to NOR task performance has not been established. Here, we investigated NOR task performance in male Lister Hooded rats following regional neural disinhibition or functional inhibition, using intra-cerebral microinfusion of the GABA-A receptor antagonist picrotoxin or agonist muscimol, respectively. Our infusion targets were the medial prefrontal cortex (mPFC), dorsal hippocampus and ventral hippocampus. Using a within-subjects design, we compared NOR task performance (1-min retention delay) following bilateral regional saline, picrotoxin or muscimol infusions made before the acquisition phase. In mPFC, neither functional inhibition nor neural disinhibition affected object recognition memory. However, in both dorsal and ventral hippocampus, neural disinhibition impaired NOR relative to saline control, mainly by reducing novel object exploration time. In addition, functional inhibition of dorsal hippocampus impaired NOR, whereas ventral hippocampal functional inhibition tended to reduce novel object exploration at the highest dose used (alongside substantial non-specific behavioural effects). Overall, our data suggest that hippocampal, but not prefrontal, GABAergic inhibition contributes to NOR at a 1-min retention delay. Moreover, such NOR performance likely requires balanced neural activity in the dorsal hippocampus, with both too little and too much dorsal hippocampal activity impairing NOR memory. Our findings support that the NOR task can be used to investigate hippocampal GABAergic dysfunction in rodent models.

**Significance statement:** Impaired GABAergic neural inhibition in the prefrontal cortex and hippocampus has emerged as a key neuropathological feature of cognitive disorders. The novel object recognition (NOR) task is used widely in rodent models to investigate cognitive impairments relevant to cognitive disorders. However, the role of hippocampal and prefrontal GABAergic inhibition in NOR is unclear, limiting interpretations as to how NOR deficits in rodent models may relate to this key pathological feature of many cognitive disorders. Here, we show that impaired hippocampal GABAergic inhibition impairs NOR in rats, whereas prefrontal GABAergic inhibition is not required. Thus, the NOR task may be used to investigate hippocampal GABAergic dysfunction in rodent models.

## 1 Introduction

Neural disinhibition, or reduced GABAergic inhibition, in the prefrontal cortex and hippocampus characterises several cognitive disorders (Lewis et al., 2005; Marín, 2012; Prévot & Sibille, 2021). In particular, disinhibition has emerged as a key neuropathological feature of schizophrenia, based on human post-mortem findings of reduced prefrontal and hippocampal GABAergic markers (Benes & Berretta, 2001; Gonzalez-Burgos et al., 2010; Heckers & Konradi, 2015). Rodent studies support that prefrontal and hippocampal GABAergic disinhibition contribute to cognitive impairments relevant to schizophrenia, including deficits in spatial memory, attention and cognitive flexibility (Bast et al., 2017; Tse et al., 2015). The novel object recognition (NOR) task is used widely to investigate cognition in rodent models, and NOR deficits have been suggested to be relevant to recognition memory impairments in human brain disorders, including schizophrenia (Cadinu et al., 2018; Grayson et al., 2015; Lyon et al., 2012; Meltzer et al., 2013). However, whether prefrontal or hippocampal GABAergic inhibition contributes to NOR memory is not clear.

A key brain region for standard (single-item) NOR is the perirhinal cortex, whereas hippocampus and medial prefrontal cortex (mPFC) are typically considered less important (Warburton & Brown, 2015). Functional inhibition and lesion studies in rodents suggest that the mPFC is not required for standard NOR at short delays (Morici et al., 2015; Nelson et al., 2011; Warburton & Brown, 2015), although there is evidence that mPFC does contribute to NOR memory consolidation or retrieval at longer (24-h) retention delays (Chao et al., 2020, 2022). Hippocampal lesion and functional inhibition studies have reported both impaired and intact NOR, which was suggested to reflect differences in NOR task procedures and testing environment (Chao et al., 2020, 2022; Cohen & Stackman Jr, 2015; Warburton & Brown, 2015). In particular, the hippocampus, especially dorsal hippocampus (DH), was suggested to be required for NOR at long (>10 min), but not short (5 min), retention delays (Ásgeirsdóttir et al., 2020; Cinalli Jr et al., 2020; Cohen & Stackman Jr, 2015; but see Neugebauer et al., 2018; Oliveira et al., 2010; Sawangjit et al., 2018).

Importantly, regardless of whether NOR requires the mPFC or hippocampus, neural disinhibition of these regions may affect NOR because regional disinhibition may disrupt processing in projection sites (Bast et al., 2017) critical for NOR. For example, the mPFC projects to the perirhinal cortex (Deacon et al., 1983) and the hippocampus also projects to regions required for NOR, including perirhinal and entorhinal cortex (Chao et al., 2022). The effect of mPFC disinhibition on NOR has, to our knowledge, not been investigated. In DH, disinhibition by infusion of the GABA-A receptor antagonist bicuculline impaired NOR at 1-min (Riordan et al., 2018) and 24-h (Kim et al., 2014) retention delays, whereas ventral hippocampus (VH) bicuculline infusion did not impair NOR at a 1-min delay (Neugebauer et al., 2018).

Here, we examined the impact of functional inhibition and neural disinhibition, induced by microinfusion of the GABA-A receptor agonist muscimol or antagonist picrotoxin, respectively, in the mPFC (experiment 1), DH (experiment 2) or VH (experiment 3) on NOR at a 1-min retention delay. Our previous in vivo electrophysiological studies confirmed that such regional muscimol/picrotoxin infusion induces neural changes consistent with inhibition/disinhibition, including reduced/enhanced burst firing (McGarrity et al., 2017; Pezze et al., 2014). The mPFC plays a limited role in standard NOR (Warburton & Brown, 2015) and, although evidence regarding hippocampal requirement for NOR is mixed, the weight of evidence suggests that the hippocampus is not required at a 1-min retention delay (Cohen & Stackman Jr, 2015). Therefore, we hypothesised that functional inhibition of mPFC, DH or VH would not impair NOR at a 1-min retention delay. However, because neural disinhibition in these regions may disrupt processing in their projection targets that are required for NOR, we hypothesised that neural disinhibition in all three regions would impair NOR.

## 2 Materials and methods

### 2.1 Rats

Three cohorts of 16 male Lister Hooded rats (experiment 1: Charles River, UK; experiments 2 and 3: ENVIGO, Harlan, UK) weighing 280-340 g and approximately 8-10 weeks old at surgery were used. For final sample sizes, sample size justification and exclusion criteria, see section 2.7. Rats were housed in groups of four in individually ventilated ‘double decker’ cages (GR1800; 462 x 403 x 404 mm; Techniplast, UK) under temperature (21 ± 1.5 °C) and humidity (50 ± 8 %) controlled conditions, and on an alternating 12-h light-dark cycle (lights on at 07:00 h) (Bio-Support Unit, University of Nottingham, UK). All procedures were carried out in the light phase, and rats had *ad libitum* access to food and water throughout the study. Rats were habituated to handling by the experimenter prior to any procedures. Procedures were conducted in accordance with the UK Animals (Scientific Procedures) Act 1986, approved by the University of Nottingham’s Animal Welfare and Ethical Review Board (AWERB) and run under the authority of Home Office project license 30/3357 and PP1257468. For the reporting of our studies, we followed ARRIVE guidelines (Percie du Sert et al., 2020).

### 2.2 Implantation of guide cannulae into medial prefrontal cortex, dorsal or ventral hippocampus

Guide cannulae were implanted above the target infusion sites in the mPFC (experiment 1), DH (experiment 2) or VH (experiment 3) (Fig. 1A), using methods similar to our previous studies (McGarrity et al., 2017; Pezze et al., 2014; Williams et al., 2022). Rats were anaesthetized using isoflurane delivered in medical oxygen (induction: 3%; maintenance: 1–3%; flow rate: 1 L/min) and prepared for surgery by shaving and disinfecting the scalp, and administering a perioperative analgesic (Rimadyl, Small Animal Solution, Zoetis, UK; 5 mg/kg, sub-cutaneous (s.c.) or Metacam, Boehringer Ingelheim; 1 mg/kg, s.c.). During the surgery, rats were secured in a stereotaxic frame. Local anaesthetic (EMLA cream 5%; lidocaine/prilocaine, Aspen, UK) was applied to the ear bars to minimise discomfort, and a topical ocular lubricant (Lubrithal, Dechra, UK) was used to protect the eyes. The scalp was incised to expose the skull, and bregma and lambda were aligned horizontally. Infusion guide cannulae (stainless steel, 26-gauge, Plastic Ones, Bilaney, UK), with stylets (stainless steel, 33-gauge, Plastics One, Bilaney, UK) inserted to prevent occlusion, were bilaterally implanted through small holes drilled in the skull. Insertion depths for guide cannulae were 0.5 mm dorsal to the infusion site, as the injectors protruded 0.5 mm from the tip of the guides. For the mPFC, we used a double guide cannula (“mouse” model C235GS-5-1.2), aimed at: AP +3.0, ML ± 0.6, DV −3.5 mm (from skull), as in Pezze et al. (2014). For the DH, we also used a double cannula (model C235G-3.0-SPC), aimed at: AP −3.0, ML ± 1.5, DV −3.5 mm (from dura); coordinates were adapted from previous studies in Wistar rats (Zhang et al., 2002a, 2002b, 2014), and based on pilot studies in Lister Hooded rats (T. Bast, unpublished data). For the VH, we implanted two separate guide cannulae (model C315G/SPC), each aimed at: AP −5.2, ML ± 4.8, DV −6.5 mm (from dura), as in Williams et al. (2022) and McGarrity et al. (2017). A stabilising pedestal was built around the cannulae using dental acrylic (Simplex Rapid, Kemdent, UK), which was anchored by four stainless-steel screws in the skull. The scalp incision was sutured around the pedestal and the rat was injected with saline (1 ml, intraperitoneal (i.p.)) to reduce risk of dehydration. The breathing rate was monitored throughout the surgery and kept at 40-60 breaths/min by adjusting the isoflurane concentration accordingly. After surgery, rats were allowed at least 5 days of recovery before testing, during which the rats were checked daily and habituated to the manual restraint necessary for the drug microinfusions. Rats also received a daily injection of prophylactic antibiotics (Synulox, Zoetis, UK; 140 mg amoxicillin, 35 mg clavulanic acid/ml; 0.2 ml/kg, s.c.), starting on the day of surgery until the end of the study.

**Fig. 1.**
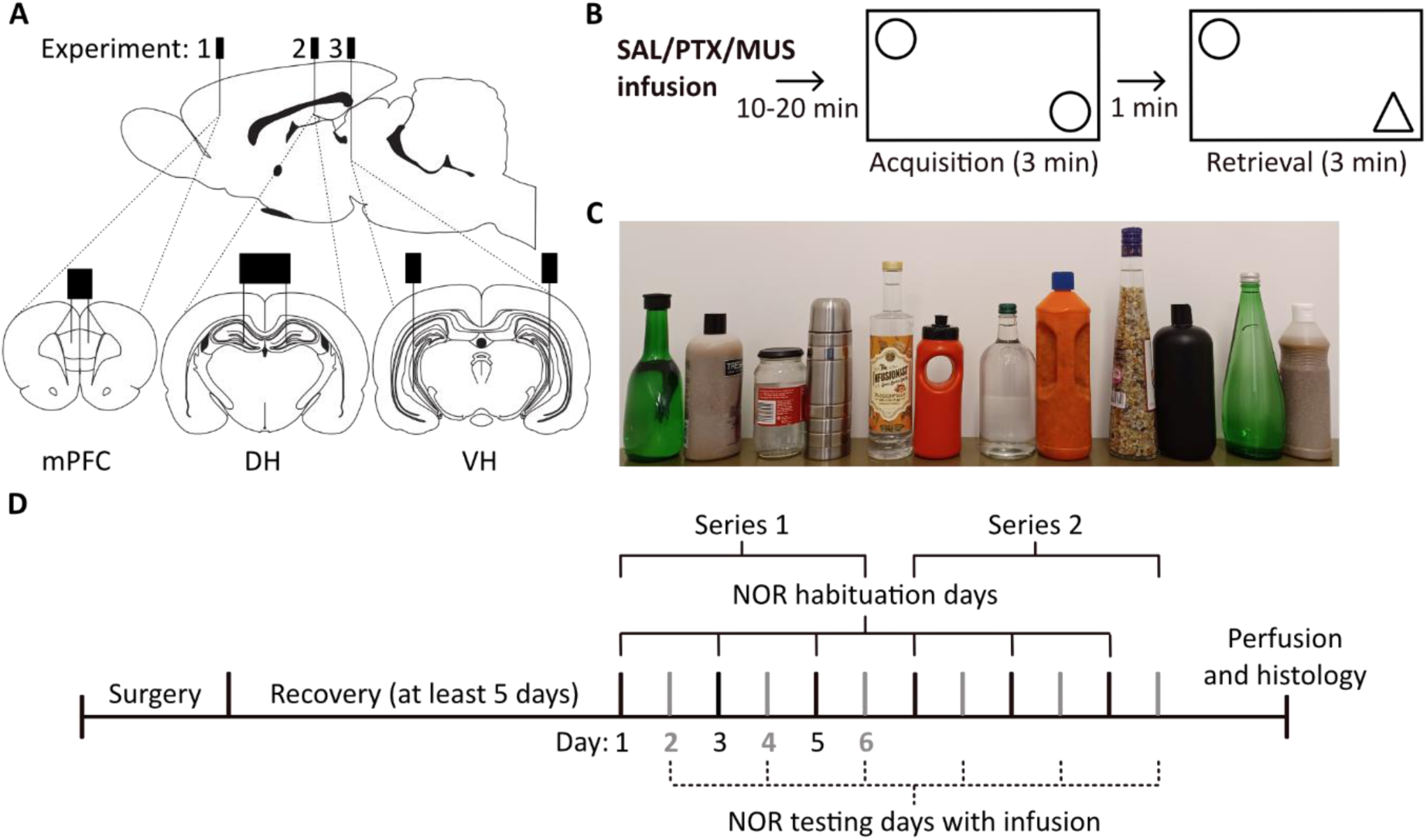
Overview of experiments to study impact of functional inhibition and disinhibition in the medial prefrontal cortex, dorsal and ventral hippocampus on novel object recognition (NOR). **A.** Target infusion sites in the medial prefrontal cortex (mPFC, experiment 1), dorsal hippocampus (DH, experiment 2), and ventral hippocampus (VH, experiment 3), adapted from Paxinos & Watson (2006). **B.** NOR testing started 10-20 min following intracerebral microinfusions of either saline (SAL), picrotoxin (PTX) or muscimol (MUS), and consisted of a 3-min acquisition and retrieval phase, separated by a 1-min retention delay. **C.** Objects used in NOR testing arranged in the pairs used as familiar and novel. **D.** Timeline of experiments. Following surgical implantation of guide cannulae and recovery, the impact of intracerebral drug infusions on NOR performance was tested using a within-subjects design, with two series of testing. Each series consisted of 6 days: NOR habituation (1 h) on days 1, 3 and 5, and infusion (saline, picrotoxin or muscimol) followed by NOR testing on days 2, 4 and 6. The order of infusion conditions was counterbalanced according to a Latin square design.

### 2.3 Microinfusion procedure and gross behavioural effects

The GABA-A receptor agonist muscimol (experiment 1: 62.5 ng/0.5 µl; experiment 2 and 3: 500 ng/0.5 µl) and antagonist picrotoxin (experiment 1: 300 ng/0.5 µl; experiment 2 and 3: 150 ng/0.5 µl) (Sigma-Aldrich, UK) were dissolved in saline. Doses used in experiment 1 were based on previous studies involving mPFC microinfusions (Pezze et al., 2014). For experiments 2 and 3, the picrotoxin dose was based on previous findings that, at 150 ng/0.5 µl/side, picrotoxin infusions into the DH or VH resulted in moderate increases in locomotor activity, without inducing seizure-related behavioural or seizure-related electrophysiological changes (Bast et al., 2001; McGarrity et al., 2017; S. McGarrity & T. Bast, unpublished findings). The muscimol dose (500 ng/0.5 µl) was based on previous studies investigating functional inhibition of the DH or VH on various behavioural tasks (Bast et al., 2001b; de Lima et al., 2006; McDonald et al., 2010; Oliveira et al., 2010; Sawangjit et al., 2018; Zhang et al., 2014). In experiment 3, the dose of picrotoxin and muscimol was reduced after the first series of testing to 100 ng/0.5 µl and 250 ng/0.5 µl, respectively, due to adverse behavioural drug effects (see below). The experimenter was blinded to the drug infusion conditions at the start of testing. However, in practice, blinding was difficult to maintain due to the presence of some behavioural drug effects that were evident from visual inspection of the rats.

Rats were gently restrained throughout the microinfusion procedure. Stylets were removed from the guide cannulae and replaced with injectors (33 gauge; Plastic Ones, Bilaney, UK), which protruded 0.5 mm below the guides. Each injector was connected to an SGE micro-syringe (5 µL; World Precision Instruments, UK) mounted on a microinfusion pump (SP200IZ syringe pump, World Precision Instruments, UK) by polyethylene tubing (PE50, Bilaney Consultants, UK). An air bubble was included in the tubing, and movement of the bubble was used to verify successful infusion of the drug into the brain. A volume of 0.5 µl/side of saline (0.9%), muscimol or picrotoxin was bilaterally infused over 1 min. The injector was kept in place for a further 1 min to allow for tissue absorption of the drug bolus. The injectors were then removed, and the stylets replaced. After the infusion, rats were placed individually into holding boxes for 10-20 min before NOR testing began (see section 2.7 for timing details). Following infusions, rats were visually inspected for any gross behavioural infusion effects.

Medial PFC infusions did not result in any behavioural changes observable by visual inspection, whereas hippocampal infusion of picrotoxin or muscimol resulted in gross behavioural changes in some rats. The gross behavioural effects of hippocampal infusions reported here are for all rats that received either a muscimol or picrotoxin infusion, regardless of whether they were included in the final NOR analysis. Following VH picrotoxin infusion (Table 1), most rats showed wet dog shakes and some other potentially seizure-related behaviours, including facial twitching and wild running, which can often be observed before full motor seizures (Lüttjohann et al., 2009; Racine, 1972; Williams et al., 2022). However, only 2 out of 15 rats infused with picrotoxin in series 1 (150 ng/side) and 1 out of 13 rats infused with picrotoxin in series 2 (100 ng/side) showed full behavioural seizures with loss of postural control. These effects were observed within 5-30 min of picrotoxin infusion. In our earlier studies using VH picrotoxin infusions of up to 150 ng/side, we did not observe behavioural seizures or any of the potentially seizure-related behavioural effects in Wistar or Lister Hooded rats (Bast et al., 2001a; McGarrity et al., 2017). In addition, in vivo electrophysiological recordings also revealed no seizure-like neural activity (McGarrity et al., 2017). However, our more recent study using VH picrotoxin (150 ng/side) revealed behavioural seizure-related effects (Williams et al., 2022). We suggested that the stress due to water restriction and fear conditioning (conditioned lick suppression test) in this previous study contributed to the higher incidence of behavioural seizure-related effects (Williams et al., 2022). However, the current study suggests that VH picrotoxin infusions at a dose of 150 ng/side can cause behavioural seizures and seizure-related effects in a substantial number of rats, even without increased stress due to aversive stimuli. Dorsal hippocampal infusion of picrotoxin (150 ng/side) caused potentially seizure-related behaviours in some rats; wet dog shakes were observed in 3 out of 16 rats, with one of those three rats also showing clonic limb movement, although we did not observe any full motor seizures.

**Table 1.** Gross behavioural effects of ventral hippocampal picrotoxin and muscimol infusions as revealed by visual inspection. Gross behavioural effects are reported for all rats that received either picrotoxin (PTX) or muscimol (MUS) infusion, regardless of whether they were included in the final analysis of NOR data. Rats were excluded from the NOR data analysis if they showed a seizure with loss of postural control or did not reach at least 5 s exploration time due to infusion-induced motor impairments. Numbers indicate rats where each behaviour was observed in either series 1 (PTX, 150 ng/0.5 µl/side, n = 15; MUS, 500 ng/0.5 µl/side, n = 16) or series 2 (PTX, 100 ng/0.5 µl/side, n = 13; MUS, 250 ng/0.5 µl/side, n = 13). None of these gross behavioural effects were observed following saline infusion.

| Observation | Series 1 PTX<br>(150 ng/side) | Series 2 PTX<br>(100 ng/side) | Series 1 MUS<br>(500 ng/side) | Series 2 MUS<br>(250 ng/side) |
| --- | --- | --- | --- | --- |
| Facial twitching | 1 | 0 | 0 | 0 |
| Wet dog shakes | 13 | 10 | 0 | 0 |
| Wild running | 2 | 1 | 0 | 0 |
| Clonic limb movement | 3 | 1 | 0 | 0 |
| Seizure with loss of postural control | 2 | 1 | 0 | 0 |
| Marked hypoactivity | 0 | 0 | 3 | 0 |
| Reversing | 0 | 0 | 3 | 0 |
| Oral tendencies | 0 | 0 | 4 | 0 |
| Agonistic behaviour in cage | 0 | 0 | 2 | 3 |

Following VH muscimol infusion (500 ng/side), we also observed some gross behavioural effects in some rats (Table 1). In three rats, VH muscimol infusion resulted in marked hypoactivity. During the NOR test sessions, these three rats often moved backwards or ‘reversed’ into the corners of the NOR arena, and showed oral tendencies, such as putting their tail in their mouth and/or licking the arena and objects. Such oral tendencies were also evident in these rats on return to their home cage following task completion. In two other rats, agonistic behaviours were observed when they were returned to the home cage; the rats would face each other standing on their hind legs, showing their teeth and occasionally hissing. This behaviour was still present 4 h following infusion but subsided within 24 h. Following VH infusions of the lower dose of muscimol (250 ng/side), we did not observe hypoactivity or unusual behaviour during the NOR sessions, but some agonistic behaviours were still observed in three rats on return to the home cage. In our previous studies using VH infusions of 500 ng/side and 1 µg/side of muscimol, we observed marked hypoactivity, but not any of the other behaviours (Bast et al., 2001b; Zhang et al., 2014). Following DH muscimol (500 ng/side) infusion, one rat showed hypoactivity and compulsive chewing and licking behaviours. The hypoactivity observed in this one rat following DH muscimol contrasts with the overall hyperactivity observed in the other rats (see section 3.2.3), and with our previous findings (Bast & Feldon, 2003).

### 2.4 NOR testing

We used a standard NOR task involving single item recognition (Ennaceur & Delacour, 1988). The rat is placed in an open field arena and allowed to explore two novel, but identical, objects for the acquisition phase and, after a retention delay, the rat is returned to the arena which now contains an identical copy of the acquisition phase object (the ‘familiar’ object) and a novel object (Fig. 1B). Due to rodents’ innate preference for novelty, rodents with intact NOR memory preferentially explore the novel object over the familiar one at test (Ennaceur & Delacour, 1988). The NOR procedure used here was adapted from previous studies (Gonçalves et al., 2023; Pezze et al., 2015) and was suitable for repeated testing of the same rat, as required for within-subjects testing of the intracerebral infusion effects.

Four rectangular NOR arenas (38 cm x 40 cm x 54 cm with a Perspex lid) were used where the brightness in each arena was 30-40 lux. An overhead camera (HD Everio, GZ-EX515BEK, JVC, UK) recorded rats’ behaviour for subsequent analyses. The objects used were mainly bottles or jars filled with sand and/or water (Fig. 1C). Object pairs (one object used as novel and one as familiar) were allocated such that they consisted of different shapes, colours, sizes and materials. The object chosen to be novel was counterbalanced across all infusion conditions, as was the position of the novel object in the arena (left or right). Time spent exploring an object was defined as interacting with the object (e.g., sniffing) and/or directing the nose towards the object from an estimated distance of < 1 cm. If the rat was in contact with the object but not facing it (e.g., standing/sitting on the object or leaning against it), this was not counted as object exploration (Ennaceur & Delacour, 1988). Rats were always placed in the arena facing the arena wall at the same location. The arena and objects were cleaned with water containing ethanol (20% v/v) before each trial to reduce olfactory cues left by the rats.

On the day before each NOR testing day, rats were placed individually into the empty arena for 1 h for habituation. On the testing day, rats were re-acclimatised to the empty arena for 3 min, before receiving one of three bilateral infusions: saline, picrotoxin or muscimol. Within 10-20 min after the infusion, rats were placed into the arena with two identical objects for the acquisition phase and allowed to explore for 3 min. After a 1-min retention delay, rats were returned to the arena, which now contained a copy of the familiar object and a novel object for the 3-min retrieval phase (Fig. 1B). Object exploration was scored from the video recordings using *‘*The Novel Object Timer’ (created by Jack Rivers-Auty, https://jackrrivers.com/program/; Gigg et al., 2020) with the scorer blind to the infusion condition and object type (novel or familiar).

### 2.5 Measurement of locomotor activity using line crossings during NOR testing

For experiments 2 and 3, the NOR arena floor was demarcated into 12 equal sectors measuring 13 cm x 14 cm to obtain a measure of locomotor activity alongside NOR testing (locomotor measurements were not obtained for experiment 1, as this change was made after completion of the experiment). The number of line crossings (or sectors crossed) were manually counted from the video recordings, with the scorer blind to the infusion condition. Line crossings were defined as instances when a rat’s front two legs and shoulders crossed a line into another sector.

### 2.6 Verification of cannula placements

At the end of the study, rats were overdosed with injectable anaesthetic (sodium pentobarbital, 200 mg/ml, approximately 1 ml, i.p., Dolethal, Vetoquinol, UK) and perfused transcardially with saline (0.9%) followed by paraformaldehyde (4% in saline). Brains were then removed and stored in paraformaldehyde (4% in saline) before they were cut into 80 µm coronal sections using a vibratome. Sections containing the relevant brain regions were mounted on microscope slides and injection sites identified using a light microscope were mapped onto coronal sections of a rat brain atlas (Paxinos & Watson, 1998). For each experiment, the sections of some rats were cresyl-violet stained for presentation purposes.

### 2.7 Experimental design and sample sizes

NOR performance following the different intracerebral drug microinfusions was compared using a within-subjects design, with two series of testing for each experiment (see Fig. 1D for timeline). Each series was run over 6 d, with habituation (1 h in empty arena) on days 1, 3 and 5, and NOR testing following saline, muscimol or picrotoxin infusion on days 2, 4 and 6, with the order of infusions counterbalanced according to a Latin square design. Series 2 of testing began one day after the end of series 1. For convenience, the four rats from the same cage were tested simultaneously on the NOR task. Rats in each cage were infused in batches of two pairs, by two experimenters: one pair of rats were infused, followed by the second pair. NOR testing started 10 min after the last rat had been infused. In practice, this meant that the delay between completion of the infusion and start of NOR testing was 10-20 min. Timings were based on previous electrophysiological measurements, where the peak effect of picrotoxin on neuronal firing in the mPFC and VH was seen between 10-30 min following infusion (McGarrity et al., 2017; Pezze et al., 2014). In experiments 1 (mPFC) and 2 (DH), the doses of picrotoxin and muscimol were the same across the two series of testing. In experiment 3 (VH), the second series was run with lower doses of picrotoxin and muscimol, based on gross behavioural effects caused by the higher doses during series 1 (see section 2.3).

All three experiments started with n = 16 rats, and we were aiming for a final sample size of n = 12-16 rats for our statistical analysis. This sample size would give a power of > 80% to detect effect sizes of Cohen’s d = 1 for differences between infusion conditions (using within-subjects pairwise comparisons, two-tailed, with a significance threshold of *p* < 0.05; G*Power; Faul et al., 2007). Rats were excluded from the NOR analysis if: an object fell over during the trial; they did not explore the objects for at least 5 s in both the acquisition and retrieval phase; they showed convulsive behavioural seizures; or if histology showed that cannulae were not located in the target brain region.

In experiment 1 (mPFC), n = 13 rats were included in the analysis; two rats were excluded due to an object falling over during NOR testing, and one rat was excluded due to the infusion cannula tips being positioned too anteriorly. In experiment 2 (DH), n = 13 rats were included in the analysis; two rats were excluded as they did not reach at least 5 s of object exploration time (one following picrotoxin infusion and one following muscimol infusion), and one rat was excluded due to objects falling over. In experiment 3 (VH), four rats were excluded from analysis of series 1 and 2. In series 1, two rats were excluded as they showed behavioural seizures following picrotoxin infusion (150 ng/side). Two more rats were excluded as they did not explore the objects for more than 5 s following muscimol infusion (500 ng/side). After completion of series 1 testing, one rat was culled due to illness. Despite reducing the doses for series 2, one rat had to be excluded from series 2 analysis because it showed a convulsive seizure following picrotoxin (100 ng/side) infusion. Overall, this gave a final sample size of n = 12 for series 1, and n = 12 for series 2, of VH drug infusions.

### 2.8 Statistical analysis

In experiments 1 and 2, which used the same doses across the two series of testing, object exploration times were analysed using a three-way repeated measures ANOVA with series (1 or 2), infusion condition (saline, picrotoxin, muscimol) and object (left versus right for acquisition-phase times and familiar versus novel for retrieval-phase times) as within-subjects factors. In experiment 3, which used different doses across the two series of testing, the data from each series were analysed separately, using a two-way ANOVA with infusion condition and object as within-subjects factors. If the ANOVA revealed a significant interaction between infusion condition and object, the simple main effect of infusion condition on familiar and novel object exploration times was examined separately. Significant main effects were examined further by post-hoc pairwise comparisons using Fisher’s LSD test (Levin et al., 1994). Paired t-tests were used for planned comparisons of familiar versus novel exploration time to test for significant NOR memory in all infusion conditions. Discrimination index (DI; time spent exploring the novel object - time spent exploring the familiar object/total time spent exploring both objects) was also used as a measure of NOR. DI values range from −1 to +1, with a higher positive value indicating greater preference for the novel object and DI = 0 indicating chance exploration (Cohen et al., 2013; Kim et al., 2014). The DI was analysed by ANOVA with infusion condition and series as within-subjects factors (experiments 1 and 2), or infusion condition as a within-subjects factor (experiment 3), and any main effects were examined further using Fisher’s LSD test. One-sample t-tests were used for planned comparisons of the DI to chance exploration (DI = 0). In experiments 2 and 3, the number of line crossings was used as a measure of locomotor activity. Line crossings were analysed using ANOVA with infusion condition and series as within-subjects factors (experiments 2), or infusion condition as a within-subjects factor (experiment 3), and any main effects examined further using Fisher’s LSD test. Graphs were generated using GraphPad Prism (version 9) and statistical tests were performed using either JASP (version 0.16.2) or SPSS (version 25) software, with *p* values of < 0.05 considered to indicate statistical significance. Data were also checked for sphericity and, if the assumption of sphericity was violated, a Greenhouse-Geisser correction of the degrees of freedom was applied. All data used for the analyses and figures presented in the present paper are shared in the supplementary material.

## 3 Results

### 3.1 Experiment 1: Medial prefrontal cortex

#### 3.1.1 Infusion sites in the medial prefrontal cortex

All rats included in the NOR analysis had infusion cannula tips located in the mPFC, within a volume that corresponded to approximately 2.7–4.2 mm anterior to bregma in the atlas by Paxinos and Watson (1998) (Fig. 2A and B).

**Fig. 2.**
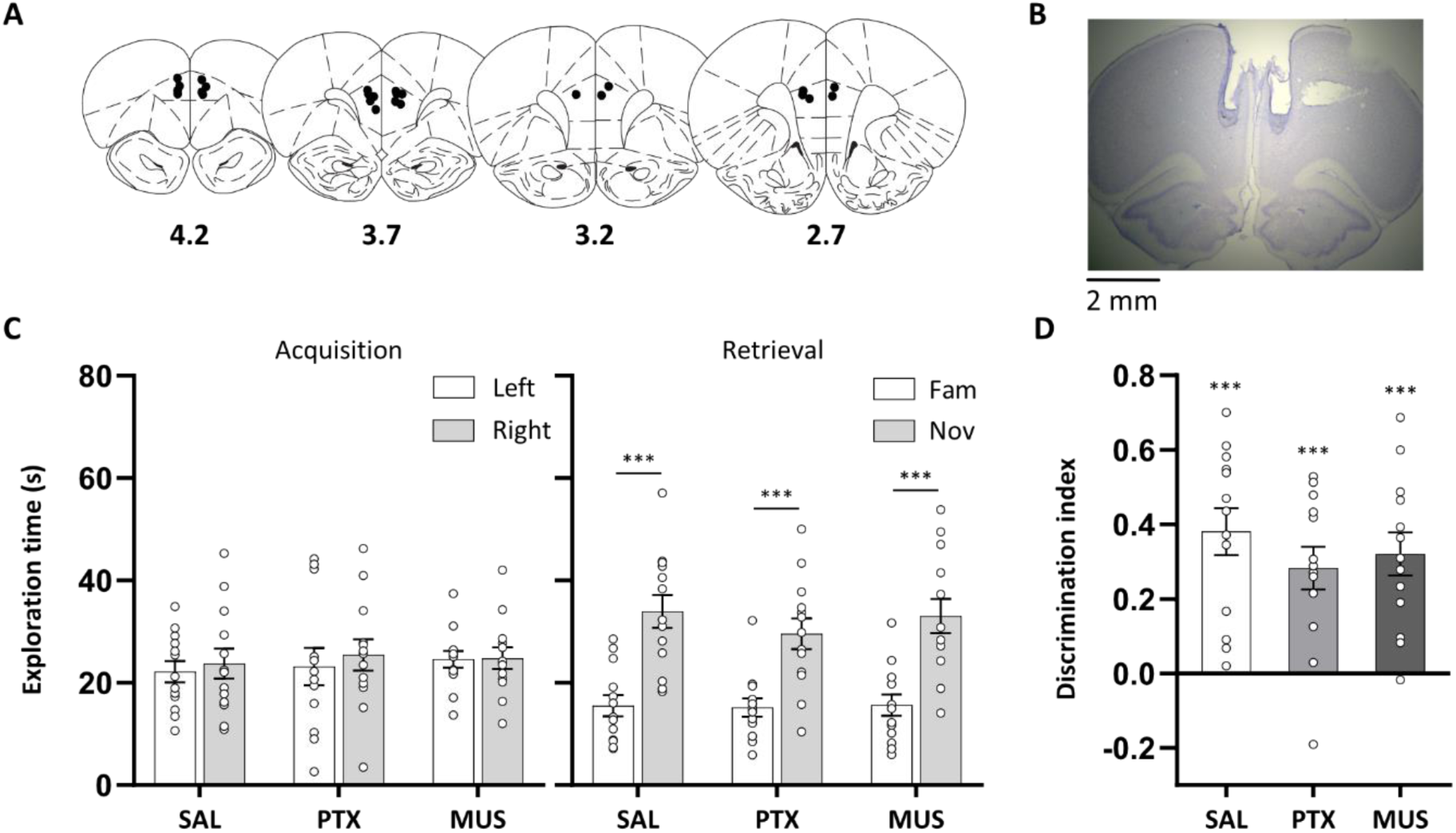
Medial prefrontal cortical disinhibition and functional inhibition did not affect novel object recognition. **A.** Approximate locations of infusion cannula tips (black dots) in the medial prefrontal cortex shown on coronal plates adapted from the atlas by Paxinos & Watson (1998), numbers indicate distance from bregma (mm). **B.** Example cresyl violet-stained coronal section showing guide cannula tracks and infusion sites in the medial prefrontal cortex. **C.** Exploration time of objects during the acquisition and retrieval phase following prefrontal disinhibition or functional inhibition, by microinfusion of picrotoxin (PTX, 300 ng/side) or muscimol (MUS, 62.5 ng/side), respectively, or following saline (SAL) control infusions. During the retrieval phase, rats across all three infusion conditions spent more time exploring the novel compared to the familiar object with no differences between infusion conditions. **D.** Discrimination index confirmed that rats in all infusion conditions showed similar novel object preference. Asterisks indicate significant novel object preference (paired or one-sample t-tests) (*p* < 0.001). Data are shown as mean (± SEM) with individual values plotted, n = 13, within-subjects design.

#### 3.1.2 Medial prefrontal cortical disinhibition and functional inhibition did not affect NOR

During acquisition, rats in all infusion conditions spent a similar amount of time exploring the two identical objects (left and right) (Fig. 2C), with ANOVA revealing no main effect of object, infusion condition or any interaction involving these factors (*F* < 1). In the retrieval phase, rats across all infusion conditions demonstrated NOR memory, with increased exploration of the novel compared to the familiar object (Fig. 2C). Planned pairwise t-tests confirmed that, in all three infusion conditions, rats spent more time exploring the novel object (*t*_12_ > 4.47, *p* < 0.001). ANOVA also revealed a main effect of object (*F*_(1, 12)_ = 64.7, *p* < 0.001), with no main effect of infusion condition or interaction (*F*_(2, 24)_ < 1). Analysis of the DI confirmed intact NOR, with novel object exploration significantly different from chance in all infusion conditions (*t*_12_ > 4.94, *p* < 0.001) and no main effect of infusion condition (*F*_(2, 24)_ < 1) (Fig. 2D).

### 3.2 Experiment 2: Dorsal hippocampus

#### 3.2.1 Infusion sites in the dorsal hippocampus

All infusion cannula tips were located in the DH, within a volume that corresponded to approximately 2.12–3.6 mm posterior to bregma in the atlas by Paxinos and Watson (1998) (Fig. 3A and B).

**Fig. 3.**
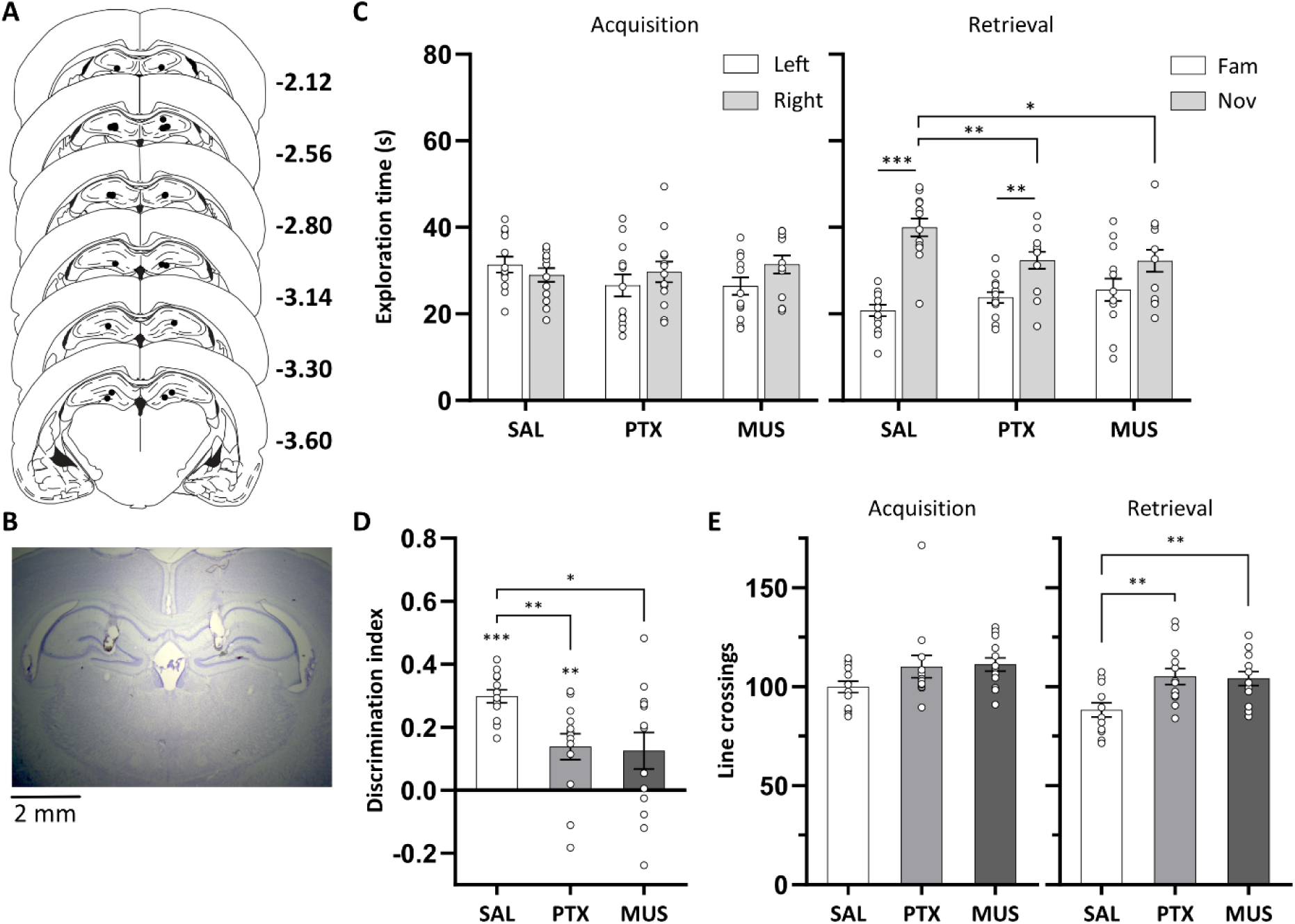
Dorsal hippocampal disinhibition and functional inhibition impaired novel object recognition. **A.** Approximate locations of infusion cannula tips (black dots) in the dorsal hippocampus shown on coronal plates adapted from the atlas by Paxinos & Watson (1998), numbers indicate distance from bregma (mm). **B.** Example cresyl violet-stained coronal section showing the end of the guide cannula tracks and, underneath, the infusion sites in the dorsal hippocampus. **C.** Exploration time of objects during the acquisition and retrieval phase following dorsal hippocampal disinhibition or functional inhibition, by microinfusion of picrotoxin (PTX, 150 ng/side) or muscimol (MUS, 500 ng/side), respectively, or following saline (SAL) control infusions. In the retrieval phase, there was a significant reduction in novel object exploration time in the picrotoxin and muscimol condition. **D.** Discrimination index was significantly reduced in the picrotoxin and muscimol condition. **E.** Picrotoxin and muscimol infusion, compared to saline, increased locomotor activity, as reflected by the number of line crossings. Asterisks indicate significant novel object preference (*p* < 0.01, paired or one-sample t-tests); or significant differences between infusion conditions (*p* < 0.05; Fisher’s LSD test following a significant main effect of infusion condition). Data are shown as mean (± SEM) with individual values plotted, n = 13, within-subjects design.

#### 3.2.2 Dorsal hippocampal disinhibition and functional inhibition impaired NOR

During acquisition, rats in all infusion conditions spent a similar amount of time exploring the two identical objects, with ANOVA revealing no main effect of infusion condition (*F*_(2,24)_ < 1) (Fig. 3C). However, ANOVA revealed a main effect of object (*F*_(1,12)_ = 9.5, *p* = 0.01), alongside a slight tendency for an interaction between object and infusion condition (*F*_(1.38,16.5)_ = 2.83, *p* = 0.102), likely due to a slight increase in exploration of objects located on the right, which was evident in the picrotoxin and muscimol conditions.

In the retrieval phase, both muscimol and picrotoxin impaired NOR (Fig. 3C). ANOVA revealed a significant interaction between infusion condition and object (*F*_(2,24)_ = 6.7, *p* = 0.005), alongside a main effect of object (*F*_(1,12)_ = 57.1, *p* < 0.001). Simple main effects analysis revealed a significant main effect of infusion condition on exploration time for novel objects (*F*_(2,24)_ = 5.16, *p* = 0.014), but not for familiar objects (*F*_(2,24)_ = 1.70, *p* = 0.204). Post hoc comparisons revealed that both picrotoxin and muscimol reduced novel object exploration time compared to saline (both *p* < 0.047), with no difference between the picrotoxin and muscimol condition (*p* = 0.966). Planned pairwise t-tests indicated that rats in the saline and picrotoxin condition explored the novel object more than the familiar object (both *t*_12_ > 3.73, *p <* 0.003), whereas rats in the muscimol condition showed a trend to preferentially explore the novel object (*t*_12_ = 1.87, *p* = 0.086).

The DI also demonstrated an NOR impairment in the picrotoxin and muscimol condition (Fig. 3D). ANOVA showed a significant main effect of infusion condition (*F*_(2,24)_ = 5.75, *p* = 0.009), reflecting a reduced DI in both the picrotoxin and muscimol condition, compared to saline (both *p* < 0.021), with no significant difference between muscimol and picrotoxin (*p* = 0.843). Planned one-sample t-tests revealed that rats in the saline and picrotoxin condition showed significant novel object preference (both *t*_12_ > 3.36, *p <* 0.006) and rats in the muscimol condition showed a trend for novel object preference (*t*_12_ = 2.17, *p* = 0.051).

#### 3.2.3 Dorsal hippocampal disinhibition and functional inhibition increased locomotor activity

Dorsal hippocampal picrotoxin and muscimol infusion increased locomotor activity during NOR testing, as indicated by an increased number of lines crossings (Fig. 3E). In the acquisition phase, picrotoxin and muscimol both tended to increase the number of line crossings, with ANOVA revealing a trend for a main effect of infusion condition (*F*_(2,24)_ = 3.12, *p* = 0.094). In the retrieval phase, ANOVA revealed a significant main effect of infusion condition (*F*_(2,24)_ = 6.42, *p* = 0.006). Post hoc comparisons revealed that both picrotoxin and muscimol increased the number of line crossings compared to saline (*p* < 0.007), with no significant difference between the picrotoxin and muscimol condition (*p* = 0.873). This increase in line crossings is consistent with our previous findings of increased open field locomotor activity following DH picrotoxin or muscimol infusion (Bast & Feldon, 2003; S. McGarrity & T. Bast, unpublished findings).

### 3.3 Experiment 3: Ventral hippocampus

#### 3.3.1 Infusion sites in the ventral hippocampus

For all rats included in the NOR analyses, infusion cannula tips were located in the VH, within a volume that corresponded to approximately 4.8–6.3 mm posterior to bregma in the atlas by Paxinos and Watson (1998) (Fig. 4A and B). In some rats, cannula infusion tips were located in the subiculum region of the VH. Nevertheless, we expect that the 0.5 µl infusion bolus would spread beyond the infusion site. If spread of the infusion volume was isotropic, the bolus would occupy a sphere with a radius of 0.5 mm centred on the infusion site. However, the spread is likely facilitated in the dorsal direction by the cannula tracks and so the drug spread is likely > 0.5 mm from the infusion sites in the dorsal direction. We expect that the drug spread would be largely limited to within the hippocampus due to the surrounding densely packed fibre bundles (Jacobs et al., 2013; Morris et al., 1989; Williams et al., 2022). Moreover, our previous multiunit recordings found that VH picrotoxin infusions, at the same coordinates as in the present study, resulted in enhanced burst firing in all VH subregions (including CA1, CA3 and dentate gyrus), with no changes in neural firing recorded outside the medial and lateral boundaries of the VH (McGarrity et al., 2017). Therefore, although some infusion cannula tips were located in the subiculum region, it is likely that the drug infusions affected all subregions of the VH, including CA1, CA3 and dentate gyrus.

**Fig. 4.**
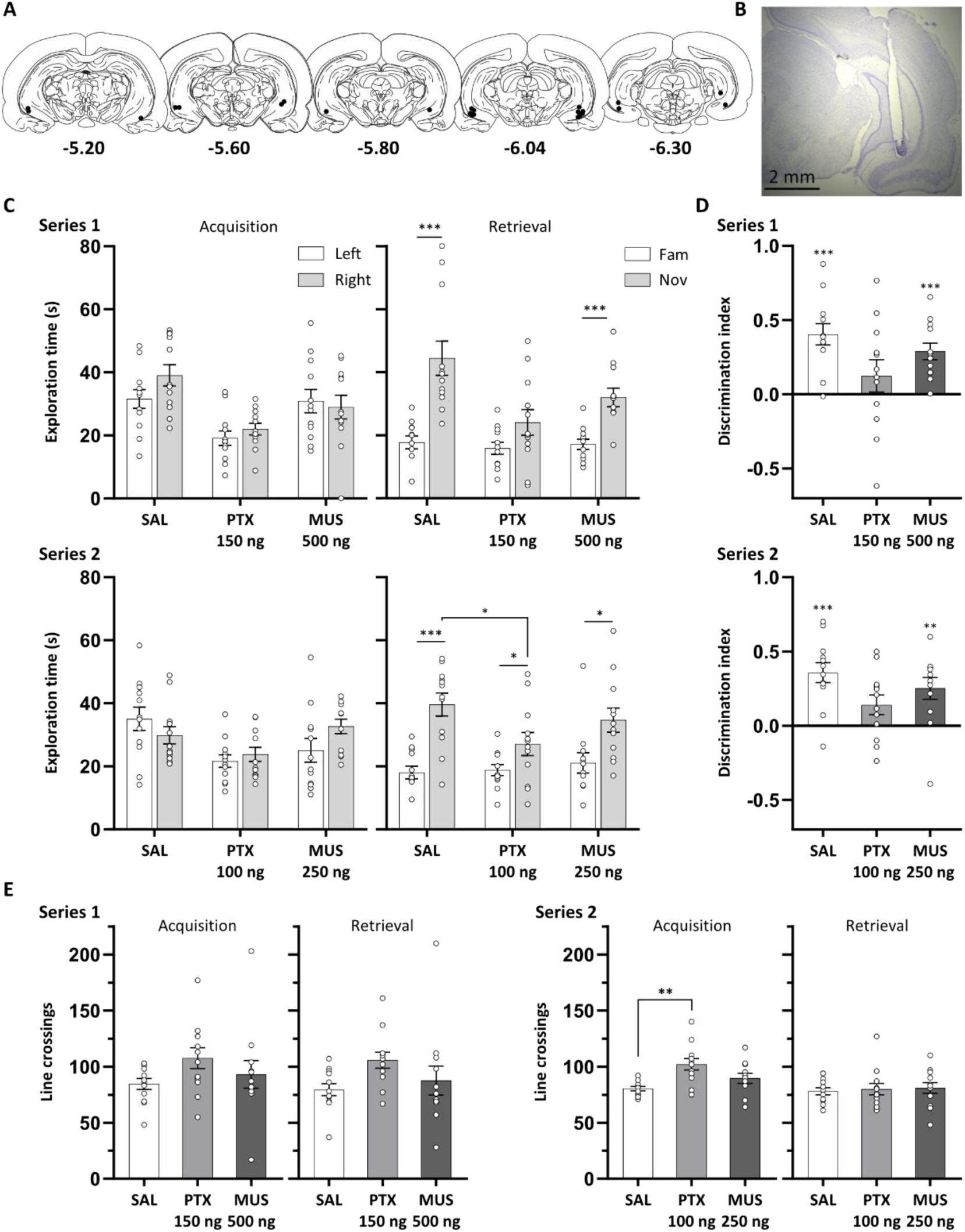
Ventral hippocampal neural disinhibition impaired novel object recognition. **A.** Approximate locations of infusion cannula tips (black dots) in the ventral hippocampus shown on coronal plates adapted from the atlas by Paxinos & Watson (1998), numbers indicate distance from bregma (mm). **B.** Example cresyl violet-stained coronal section showing a guide cannula track and, underneath, an infusion site in the ventral hippocampus. **C.** Exploration times of objects during the acquisition and retrieval phase in series 1 and 2. In series 2, novel object exploration was reduced by picrotoxin (PTX, 100 ng/side), but not muscimol (MUS, 250 ng/side), infusion compared to saline (SAL) control. **D.** Discrimination index in both series was not significantly above chance following PTX infusion. **E.** Line crossings increased following PTX infusion in series 1 and 2. Asterisks indicate significant novel object preference (*p* < 0.05, paired or one-sample t-tests); or significant differences between infusion conditions (*p* < 0.05; Fisher’s LSD test following a significant main effect of infusion condition). Data are shown as mean (± SEM) with individual values plotted, n = 12, within-subjects design.

#### 3.3.2 Ventral hippocampal disinhibition impaired NOR

During acquisition, rats in all infusion conditions explored the identical left and right objects for a similar amount of time in both series (series 1: saline, 150 ng/side picrotoxin, 500 ng/side muscimol; series 2: saline, 100 ng/side picrotoxin, 250 ng/side muscimol) (Fig. 4C), with ANOVA indicating no main effect of object for both series 1 and 2 (both *F*_(1,11)_ < 1.66, *p* > 0.224). However, picrotoxin infusions reduced overall exploration time. ANOVA revealed a main effect of infusion condition for both series (*F*_(2,22)_ > 6.41, *p* < 0.006), with post hoc pairwise comparisons indicating that this was due to a decrease in overall exploration time in the picrotoxin condition compared to the saline and muscimol condition (all p < 0.022), which did not differ (both series *p* > 0.228). In series 1, there was no interaction between object and infusion condition (*F*_(2,22)_ = 1.70, *p* = 0.206), whereas, in series 2, the object and infusion condition interaction was at trend level (*F*_(2,22)_ = 3.14, *p* = 0.063). This is likely due to rats in the saline condition numerically exploring the left object more than the right one, whereas an opposite numerical preference was shown by rats in the picrotoxin and muscimol condition.

During the retrieval phase in series 1, picrotoxin (150 ng/side) or muscimol (500 ng/side) infusion tended to impair NOR performance, demonstrated by a trend towards reduced exploration of the novel object, compared with the saline condition (Fig. 4C, series 1). ANOVA revealed a main effect of object (*F*_(1,11)_ = 64.8, *p* < 0.001) and infusion condition (*F*_(2,22)_ = 6.85, *p* = 0.005). Post hoc comparisons indicated a significant reduction in overall exploration time in the picrotoxin and muscimol condition compared to saline (both *p* < 0.024), with no significant difference between picrotoxin and muscimol (*p* = 0.163). The infusion and object interaction was at trend level (*F*_(2,22)_ = 3.15, *p* = 0.063), reflecting that novel object preference tended to be less pronounced in the picrotoxin and muscimol infusion compared to saline. Planned pairwise t-tests indicated that rats explored the novel object significantly more than the familiar object following saline and muscimol infusions (both *t*_11_ > 4.21, *p* = 0.001), and tended to show such novel object preference following picrotoxin infusion (*t*_11_ = 1.90, *p* = 0.085).

ANOVA of the DI did not reveal a significant main effect of infusion (*F*_(2,22)_ = 2.29, *p* = 0.125) (Fig. 4D, series 1). Planned one-sample t-tests demonstrated that the DI was significantly different from chance in the muscimol and saline condition (both *t*_11_ > 5.22, *p* < 0.001), but not in the picrotoxin condition (*t*_11_ = 1.13, *p* = 0.281). Overall, series 1 of the VH infusion study showed a trend for reduced novel object exploration in the picrotoxin and muscimol condition compared to saline, and planned comparisons of exploration times for novel and familiar objects, and of the DI to chance, indicated that rats in the picrotoxin condition did not preferentially explore the novel object. However, no changes were found in the DI between infusion conditions. In addition, interpretation of these findings may be limited by the reduced overall object exploration time found in the picrotoxin condition at acquisition, and in the picrotoxin and muscimol condition during retrieval.

During the retrieval phase in series 2, VH picrotoxin (100 ng/side) reduced novel object exploration compared to the saline condition, whereas muscimol (250 ng/side) did not (Fig. 4C, series 2). ANOVA revealed a significant interaction between infusion condition and object (*F*_(2,22)_ = 3.74, *p* = 0.04), alongside a main effect of object (*F*_(1,11)_ = 22.0, *p* = 0.001) and no main effect of infusion condition (*F*_(2,22)_ = 1.99, *p* = 0.160). Simple main effects analysis revealed a significant main effect of infusion condition for novel object exploration time (*F*_(2,22)_ = 4.04, *p* = 0.032), but not familiar object exploration time (*F*_(2,22)_ < 1). Post hoc comparisons revealed that VH picrotoxin infusion reduced novel object exploration time compared to saline (*p* = 0.03) and tended to reduce novel exploration time compared to muscimol (*p* = 0.062), with no difference between muscimol and saline (*p* = 0.296). Nevertheless, rats in all infusion conditions showed significant novel object preference, with planned pairwise t-tests demonstrating a significant difference between familiar and novel object exploration times in all infusion conditions (all *t*_11_ > 2.30, *p* < 0.042). ANOVA of the DI (Fig. 4D, series 2) demonstrated a trend for a main effect of infusion condition (*F*_(2,22)_ = 2.93, *p* = 0.074). Planned one-sample t-tests indicated that the DI was above chance following saline and muscimol infusion (both *t*_11_ > 3.45, *p* < 0.005) and tended to be above chance following picrotoxin (*t*_11_ = 2.1, *p* = 0.059). Overall, series 2 of the VH infusion study suggested reduced novel object exploration following picrotoxin, but not muscimol, infusion. Nevertheless, following picrotoxin infusions rats showed or tended to show significant novel object preference, along with the other infusion conditions. As for series 1, the interpretation of these findings may be limited by the reduced overall object exploration time following picrotoxin infusion during the acquisition phase.

#### 3.3.3 Ventral hippocampal disinhibition increased locomotor activity

In series 1, there was a numerical increase in line crossings in the picrotoxin condition in both the acquisition and retrieval phase, although this was not significant during either phase (*F*_(2,22)_ < 2.21, *p* > 0.133) (Fig. 4E, series 1). In series 2, during acquisition, line crossings were increased in the picrotoxin condition (Fig. 4E, series 2). ANOVA revealed a main effect of infusion (*F*_(2,22)_ = 7.07, *p* = 0.004), with post hoc comparisons indicating a significant increase following picrotoxin compared to saline infusion (*p* = 0.001), with no other differences (both *p* > 0.1). The increases in line crossings are consistent with our previous findings of increased open field locomotor activity following VH picrotoxin infusion (Bast et al., 2001a; McGarrity et al., 2017). In the retrieval phase, rats showed a similar number of line crossings regardless of infusion condition (*F*_(2,22)_ < 1).

## 4 Discussion

Our findings suggest that balanced neural activity within the hippocampus, but not mPFC, is required for NOR over 1-min retention delays. Dorsal and ventral hippocampal disinhibition by picrotoxin impaired NOR, demonstrating that hippocampal GABAergic inhibition is required for intact NOR. Dorsal hippocampal functional inhibition by muscimol also impaired NOR, whereas there was limited evidence for impairments following VH functional inhibition. In contrast, neither functional inhibition nor disinhibition of the mPFC affected NOR.

### Prefrontal functional inhibition and disinhibition did not affect NOR

Intact NOR following mPFC muscimol is consistent with a range of lesion and inactivation studies demonstrating a limited role for mPFC in standard NOR performance over retention delays of < 24 h (Chao et al., 2022; Warburton & Brown, 2015) and, specifically, with a study in rats where mPFC muscimol did not affect NOR over a 1-min delay (Neugebauer et al., 2018). Prefrontal disinhibition by picrotoxin also did not affect NOR, suggesting that mPFC GABAergic function is not required over 1-min delays. Consistent with this, optogenetic mPFC stimulation after acquisition did not affect short-term (5-min) NOR memory (Benn et al., 2016).

### Functional inhibition of dorsal hippocampus impaired NOR, whereas ventral hippocampal functional inhibition had limited effects

Ventral hippocampal muscimol slightly reduced novel object exploration, but left the DI largely unaffected, at a dose of 500 ng/side, which also reduced locomotor activity. However, VH infusion of a lower muscimol dose (250 ng/side) did not affect NOR and also did not reduce locomotor activity. Consistent with the limited impact of VH muscimol, VH optogenetic inhibition during retrieval did not affect NOR at a 10-min delay (Sun et al., 2020). In contrast, VH infusion of a relatively low dose of muscimol (50 ng/side) before acquisition impaired NOR at a 1-min delay (Neugebauer et al., 2018).

Previous lesion and temporary inactivation studies examining DH requirement for NOR reported mixed results (reviewed in Chao et al., 2020, 2022). To reconcile these mixed findings, Cohen & Stackman Jr (2015) proposed that the hippocampus is only required for NOR with delays >10 min and when a 30-s threshold of object exploration is met during acquisition. In support of this proposal, DH muscimol impaired long-term (20-min), but not short-term (5-min), NOR memory (Ásgeirsdóttir et al., 2020), and, at a 24-h delay, DH muscimol impaired NOR when sample objects were explored for 30 s, but not 10 s (Cinalli Jr et al., 2020). However, our findings do not fully support the Cohen & Stackman Jr (2015) proposal. Consistent with the suggestion that exploration times >30 s render NOR hippocampus dependent, all rats in experiment 2 explored the sample objects for >30 s, except for one (exploration was 24 s and exclusion did not affect the finding). However, contrary to the suggestion that the hippocampus is not required at delays <10 min, we found that DH muscimol impaired NOR at a 1-min delay. Hippocampal lesion studies have also demonstrated recognition memory deficits at short (≤5 min) delays in rodents (Gaskin et al., 2010; McHugh et al., 2022) and monkeys (Zola et al., 2000). In further support of DH contributions to NOR at short retention delays, optogenetic attenuation of DH CA2/CA3 to CA1 projections impaired NOR at a 5-min delay (Raam et al., 2017) and optogenetic silencing of DH adult-born dentate gyrus neurons during test impaired performance on a continuous NOR task at a <5 min delay (McHugh et al., 2022).

While our findings suggest that DH is required for NOR, overall evidence for DH requirement in NOR is mixed. The latter may reflect the use of different NOR procedures and that NOR is sensitive to task procedure with length of habituation, object exploration, and retention delay influencing performance (Cohen & Stackman Jr, 2015; Oliveira et al., 2010; Stefanko et al., 2009). Primate hippocampal lesion studies have also reported mixed effects on single item visual recognition, although a recent meta-analysis found an overall impairment (albeit smaller than that caused by perirhinal lesions) (Waters et al., 2023). Importantly, our findings are consistent with the idea that the hippocampus is important for binding a range of object features from different sensory modalities into one memory representation (Olsen et al., 2012; Squire & Zola-Morgan, 1991).

Our finding that NOR is more sensitive to DH than VH functional inhibition is consistent with the proposal that these regions support different aspects of recognition memory, with DH contributing to encoding of objects (and their context/location), whereas VH links events within a context and distinguishes different contexts (Preston & Eichenbaum, 2013). Given that the NOR deficits in the present study may partly reflect impaired novelty processing (see below), the greater sensitivity of NOR to DH functional inhibition is also consistent with a recent human study implicating the posterior hippocampus (which corresponds to rodent DH), but not anterior hippocampus (which corresponds to VH), in processing novelty (Angeli et al., 2025).

### Disinhibition of dorsal hippocampus and ventral hippocampus impaired NOR

Impaired NOR following DH disinhibition is consistent with studies demonstrating deficits following DH bicuculline infusion at 1-min (Riordan et al., 2018) and 24-h (Kim et al., 2014) retention delays. Furthermore, it is consistent with a recent study suggesting that a specific type of GABAergic interneurons in DH CA1 is important for encoding novel information underlying recognition memory (Tamboli et al., 2024). Consistent with our finding of impaired NOR following VH disinhibition by picrotoxin (100 ng/side), VH optogenetic stimulation during the retrieval phase impaired NOR at a 2-min delay (Kapanaiah et al., 2024). However, Neugebauer et al. (2018) reported intact NOR (1-min delay) following VH bicuculline infusion. In the present study, VH disinhibition reduced sample object exploration. This likely reflects that increased locomotor activity, a consistent effect of VH picrotoxin (Bast et al., 2001a; McGarrity et al., 2017), detracted from object exploration. Reduced object exploration during acquisition could result in weaker memory, impairing subsequent object recognition (compare Ainge et al., 2006). However, exploration times in experiment 3 were all above 30 s, suggesting sufficient exploration for ‘strong’ memories (Cohen & Stackman Jr, 2015).

Both DH functional inhibition and disinhibition impaired NOR, suggesting NOR requires balanced DH activity. In contrast, VH functional inhibition had limited effects, i.e. VH processing was not required, suggesting NOR impairments by VH disinhibition were due to aberrant drive of projections to other regions. Our recent metabolic imaging study revealed that, following VH disinhibition by picrotoxin, regional cerebral blood flow increased in the VH, but decreased in the DH (Williams et al., 2019), possibly reflecting VH feedforward inhibition (Sik et al., 1994). Therefore, NOR impairment by VH disinhibition could, partly, reflect DH deactivation. Moreover, VH or DH disinhibition may impair NOR due to aberrant drive of projections to extrahippocampal regions important for NOR, including perirhinal and entorhinal cortex (Chao et al., 2022).

### Hippocampal disinhibition and functional inhibition may impair novelty processing

NOR impairments in the present study mostly manifested as reduced novel object exploration, with familiar object exploration not significantly affected (although there were numerical increases). Comparison of this finding to other studies is difficult, as often only the DI is reported. If NOR is impaired, rats may explore both objects as novel (impaired familiarity detection) or both objects as familiar (impaired novelty detection). Novelty and familiarity processing were suggested to be partly independent and make distinct contributions to recognition (Kafkas & Montaldi, 2018). Consistent with the long-standing idea that the hippocampus is important for novelty detection (Kumaran & Maguire, 2007; Olsen et al., 2012; Vinogradova, 2001), it was suggested that the hippocampus contributes to novelty detection, whereas parahippocampal regions contribute to relative familiarity assessment (Kafkas & Montaldi, 2018). Therefore, NOR deficits in the present study may, partly, reflect deficits in hippocampal novelty processing (Mumby, 2001). A general impairment in novelty detection should also reduce object exploration during acquisition. Object exploration during acquisition was reduced by VH picrotoxin, although this likely resulted from locomotor changes (see above), but was not affected by the other hippocampal manipulations. Therefore, our findings may reflect that hippocampal manipulations only disrupt novelty detection of one object compared to another, which requires relatively accurate object representation, but leave intact the easier task of detecting the novelty of two objects compared to a familiar background. Alternatively, the reduced novel object exploration may reflect a false memory for the novel object, resulting in it being explored as familiar (McTighe et al., 2010), or a reduced perceived salience of the novel object, consistent with a link between hippocampal activity and salience attribution (Kätzel et al., 2020). These interpretations demonstrate the difficulty in viewing the NOR task as a pure assessment of recognition memory.

## Conclusions

Overall, NOR at a 1-min retention delay requires balanced hippocampal, but not prefrontal, activity. Object recognition memory was disrupted by reduced GABAergic inhibition in the DH and VH, but not mPFC, suggesting the NOR task can be used to study impaired hippocampal GABAergic inhibition in rodent models. Furthermore, our findings add to evidence that the DH can be required for NOR, although the factors determining whether NOR requires the hippocampus remain to be determined.

## Supporting information

Data file for Taylor et al.

## Author contributions

CT, PM, JG, MH, JN and TB designed research.

CT, JR, JL, MG, SW and RGA performed research.

CT analysed data.

CT and TB wrote the original draft of the manuscript, with all other authors subsequently contributing to revising the draft; all authors have approved the final content.

## Acknowledgements

This study and CT were supported by the Biotechnology and Biological Sciences Research Council (BBSRC) Doctoral Training Programme (DTP) CASE award at the University of Nottingham in partnership with b-neuro (grant number BB/M008770/1, project 2270910). JR and MG were supported by BBSRC DTP CASE awards in partnership with Boehringer Ingelheim; SW and RGA were supported by BBSRC DTP studentships; JL was supported by a Medical Research Council (MRC) Integrated Midlands Partnership for Biomedical Training (IMPACT) DTP studentship.

